# Impact of RNA degradation on measurements of gene expression

**DOI:** 10.1101/002261

**Authors:** Irene Gallego Romero, Athma A. Pai, Jenny Tung, Yoav Gilad

## Abstract

The use of low quality RNA samples in whole-genome gene expression profiling remains controversial. It is unclear if transcript degradation in low quality RNA samples occurs uniformly, in which case the effects of degradation can be normalized, or whether different transcripts are degraded at different rates, potentially biasing measurements of expression levels. This concern has rendered the use of low quality RNA samples in whole-genome expression profiling problematic. Yet, low quality samples are at times the sole means of addressing specific questions – e.g., samples collected in the course of fieldwork. We sought to quantify the impact of variation in RNA quality on estimates of gene expression levels based on RNA-seq data. To do so, we collected expression data from tissue samples that were allowed to decay for varying amounts of time prior to RNA extraction. The RNA samples we collected spanned the entire range of RNA Integrity Number (RIN) values (a quality metric commonly used to assess RNA quality). We observed widespread effects of RNA quality on measurements of gene expression levels, as well as a slight but significant loss of library complexity in more degraded samples. While standard normalizations failed to account for the effects of degradation, we found that a simple linear model that controls for the effects of RIN can correct for the majority of these effects. We conclude that in instances where RIN and the effect of interest are not associated, this approach can help recover biologically meaningful signals in data from degraded RNA samples.

## Introduction

Degradation of RNA transcripts by the cellular machinery is a complex and highly regulated process. In live cells and tissues, abundance of mRNA is tightly regulated, and transcripts are degraded at different rates by various mechanisms [1], partially in relation to their biological function [2–5]. In contrast, the fates of RNA transcripts in dying tissues, and the decay of isolated RNA, are not part of normal cellular physiology and therefore less likely to be tightly regulated. In spite of a few studies that addressed this issue, it remains largely unclear whether most transcript types decay at similar rates under such conditions, or whether rates of RNA decay in dying tissues are associated with transcript-specific properties.

These questions are of great importance for studies that rely on sample collection in the field or in clinical settings (both from human populations as well as from other species), where tissue samples often cannot immediately be stored in conditions that prevent RNA degradation. In these settings, extracted RNA is often partly degraded and may not faithfully represent *in vivo* gene expression levels. Sample storage in stabilizers like RNALater lessens this problem [6] but is not always a feasible approach. Differences in RNA quality and sample handling could therefore confound subsequent analyses, especially if samples subjected to different amounts of degradation are naïvely compared against each other. The degree to which this confounder affects estimates of gene expression levels is unknown.

Similarly, there is no consensus on the level of RNA decay that renders a sample unusable, or on approaches to control for the effect of *ex vivo* processes in the analysis of gene expression data. Thus, while standardized RNA quality metrics like the Degradometer [7] or the RNA Integrity Number (RIN; [8]) provide well-defined empirical methods to assess and compare sample quality, there is no widely accepted criterion for sample inclusion. For example, proposed thresholds for sample inclusion have varied between RIN values as high as 8 [9] and as low as 3.95 [10]. The recent GTEx project (http://www.broadinstitute.org/gtex/), meanwhile, reports both the number of total RNA samples they collected as well as the number of RNA samples with RIN scores higher than 6, presumably as a measure of the number of high quality samples in the study.

Broadly speaking, three approaches can be adopted to deal with RNA samples of variable quality. First, RNA samples with evidence of substantial degradation can be excluded from further study; this approach relies on establishing a cut-off value for “high quality” versus “low quality” samples, and suffers from the current lack of consensus on what this cut-off should be. It also could exclude the possibility of ever including in a study unique and hard to collect samples from remote locations. Second, if investigators are willing to assume that all transcript types decay at a similar rate, variation in gene expression estimates due to differences in RNA integrity could be accounted for by applying standard normalization procedures. Third, if different transcripts decay at different rates, and if these rates are consistent across samples for a given level of RNA degradation – e.g. a given RIN value – a model that explicitly incorporates measured, sample-specific, degradation could be applied to gene expression data to correct for the confounding effects of degradation.

To date, work on the effects of RNA decay has not provided clear guidelines with respect to these approaches. In addition, nearly all published work that focuses on RNA stability in tissues following cell death and/or sample isolation predates, or does not employ, high throughput sequencing technologies. These studies broadly suggest that both the quantity and quality of recovered RNA from tissues could be affected by acute pre-mortem stressors like pyrexia or prolonged hypoxia (11–13), and by the time to sample preservation and RNA extraction. The quantity and quality of recovered RNA are strongly dependent on the type of tissue studied [11], even when sampling from the same individual [12, 13]. These differences in yield across tissues have resulted in a wide range of recommendations for an acceptable *post-mortem* interval for extracting usable, high-quality RNA, ranging from as little as 10 minutes [14] to upwards of 48 hours [15], depending on tissue source and preservation conditions.

Similarly, studies examining changes in the relative abundance of specific transcripts as a result of *ex vivo* RNA decay have reached somewhat contradictory recommendations – although some of this conflict may be attributable to methodological differences. Studies that focused on small numbers of human genes assayed through quantitative PCR consistently report little to no effect of variation in RNA quality on gene expression estimates (6, 19–22). Conversely, microarray-based studies have repeatedly reported significant effects of variation of RNA quality on gene expression estimates, even after standard normalization approaches have been used. For example, increasing the time from tissue harvesting to RNA extraction or cryopreservation from 0 to only 40 or 60 minutes significantly affected expression profiles in roughly 70% of surveyed genes in an experiment on human colon cancer tissues [16]. Likewise, a substantial fraction of genes in peripheral blood mononuclear cells (PBMCs) appears to be very sensitive to *ex vivo* incubation [17]. Other microarray-based studies have reached similar conclusions, both in humans [11, 12, 18, 19] and other organisms [20], and have urged for caution when analyzing RNA samples with medium or low RIN scores, although the definition of an acceptable RNA quality threshold remain elusive.

To address this issue, we sequenced RNA extracted from PBMC samples that were stored unprocessed at room temperature for different time periods, up to 84 hours. Our design is aimed at mimicking sample collection in field studies. We collected RNA decay time-course data spanning almost the entire RIN quality scale, and examined relative gene-specific degradation rates through RNA sequencing. Due to the high sensitivity and resolution of high-throughput RNA sequencing, our data provide an unprecedentedly detailed picture of the dynamics of RNA degradation in stressed, *ex vivo* cells. Based on our results, we develop specific recommendations for accounting for these effects in gene expression studies.

## Results

We extracted RNA from 32 aliquots of PBMC samples from four individuals. The PBMC samples were stored at room temperature for 0 h, 12 h, 24 h, 36 h, 48 h, 60 h, 72 h and 84 h prior to RNA extraction. As expected, time to extraction significantly affected the RNA quality (p < 10^−11^), with mean RIN = 9.3 at 0 h, and 3.8 at 84 h (supplementary table 1). Based on the RIN values we chose to focus on 20 samples from five time points (0 h, 12 h, 24 h, 48 h and 84 h) that spanned the entire scale of RNA quality. We generated poly-A-enriched RNA sequencing libraries from the 20 samples using a standard RNA sequencing library preparation protocol (see [21]). We added a spike-in of non-human control RNA to each sample, which allowed us to confirm the effects of RNA degradation on the RNA sequencing results (see methods for more details). Following sequencing, we randomly subsampled all libraries to a depth of 12,129,475 reads, the lowest number of reads/library observed in the data. We used BWA 0.6.3 to map reads, calculated RPKM, and normalized the data using a standard quantile normalization approach (e.g. as in [22]). We observed that sample RIN is associated with both the number of uniquely mapped reads (ANOVA p < 10^−3^), and the number of reads mapped to genes (p = p < 10^−3^; supplementary figure 1), with high RIN samples having greater numbers of both. Furthermore, the proportion of exogenous spike-in reads increases significantly as RIN decreases (p < 10^−10^), as expected given degradation-driven loss of intact human transcripts in poor quality samples. Sequence reads from one individual were poorly mapped, especially in the later time-points (see methods and supplementary figure 1), and so we excluded the data from all samples from this individual in subsequent analysis.

### The effect of RNA degradation on RNAseq output

Principal component analysis of our data demonstrates that much of the variation (28.9%) in gene expression levels in our study is strongly associated with RNA sample RIN scores (figure 1a; PC1 associated with RIN scores p < 10^−7^; no other PCs are significantly associated with either sample storage time or RIN score; supplementary table 2). We also observed a residual presence of inter-individual variation in the data, in spite of variable RNA quality (PCs 4 and 5; supplementary figure 2 and supplementary table 2). A correlation matrix based on the gene expression data (figure 1b) indicates that while samples of relatively high quality RNA cluster by individual, data from RNA samples that experienced high yet similar degradation levels (namely, from the same later time-points) are more correlated than data from samples from the same individual across the time-points. This pattern contrasts with the naïve expectation that gene expression differences between individuals should be the strongest signal in the data. Instead, inter-individual differences only predominate in the early stages of degradation, at the early time-points of 0 h (mean RIN = 9.3) and 12 h (mean RIN = 7.9). These observations are robust with respect to the approach used to estimate gene expression levels, and – importantly – are not explained by unequal rates of degradation occurring at different distances from the 3’ poly-A tail. For example, we found nearly identical patterns when we estimated expression levels based only on reads that map to the 1000 bp at the 3’ end of each gene (supplementary figure 3). Furthermore, these global effects of RNA degradation on the estimated gene expression levels cannot be accounted for by simply regressing out RIN scores (supplementary figure 4).

**Fig 1.**
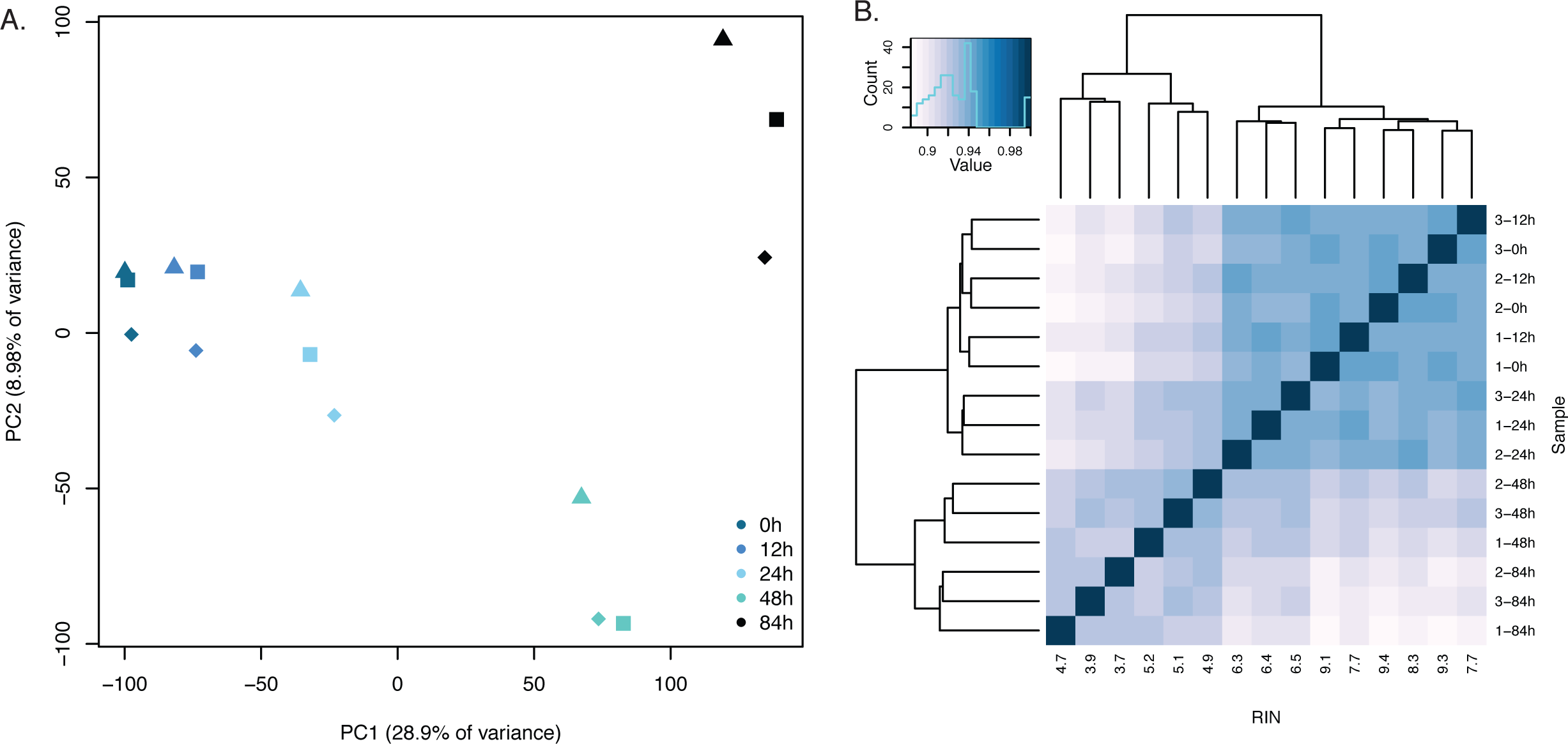
a. PCA plot of the 15 samples included in the study based on data from 29,156 genes with at least one mapped read in a single individual. Different shapes indicate different individuals. b. Spearman correlation plot of the 15 samples in the study.

The possibility of reduced sequencing library complexity is often cited as a reason to exclude RNA samples of low quality. This concern is mostly based on the observation that sequencing RNA samples of lower RNA quality results in relatively decreased proportion of mappable reads, an observation corroborated in our study (supplementary figure 1). Yet, it is unclear to what extent this property affects the ability to estimate gene expression levels in RNA samples of low quality. To assess the effects of RIN on sample complexity, we plotted the distribution of RPKM values within individuals at different time points. Our data indicate that mean RPKM increases as sample RIN decreases (p < 10^−5^). This seems counterintuitive, yet the reason is the presence of a few highly amplified genes in the samples of low RNA quality. Indeed, relative to 0 h, low RIN samples at 48 h and 84 h have an excess of low RPKM genes and a deficit of high RPKM genes, shifting the median RPKM downwards (p < 10^−4^; figure 2). We further found a positive association between the number of genes with at least one observation of RPKM >= 0.3 and RIN (p < 10^−4^). Even when we subsampled all samples to the same number of sequencing reads, we still observed a high proportion of genes with low RPKM values in RNA samples of lower quality (p < 10^−4^; supplementary figure 5). This suggests that a non-uniform effect of RNA degradation on gene expression levels results in somewhat lower complexity of the sequencing library (figure 2, supplementary figure 5). On the other hand, both within a single individual and across the whole dataset, we found that nearly all genes whose expression could be measured at 0 h are also detected as expressed throughout the entire time-course experiment, with only very few genes present in all individuals up until a given time point completely absent from the data onwards (table 1).

**Fig 2.**
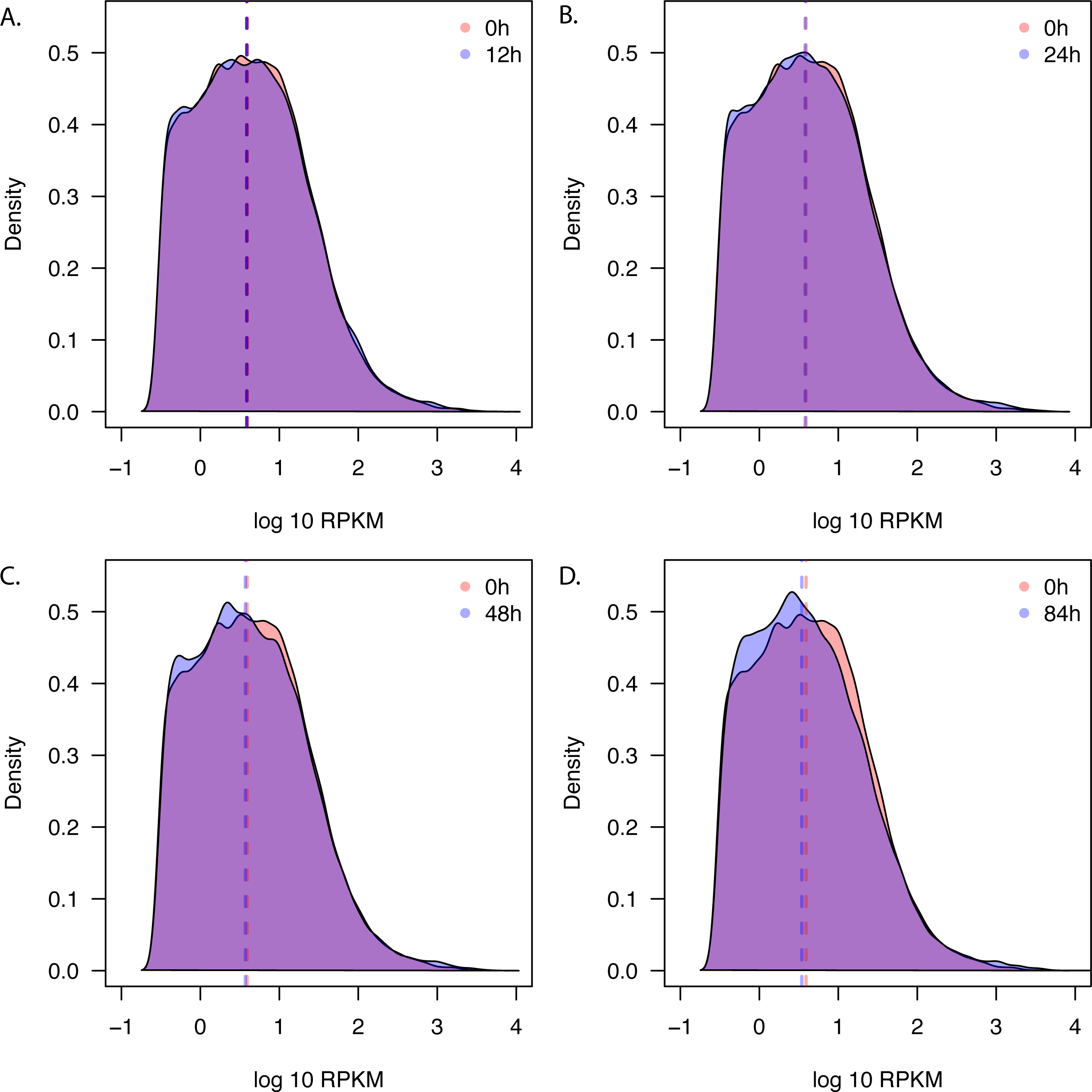
Changes in library complexity over time. Dashed lines indicate median RPKM at each time-point. a. Density plots of RPKM values amongst all three individuals at 0 h and 12 h. b. as a, but 0 h and 24 h. c. as a, but 0 h and 48 h. d. as a, but 0 h and 84 h.

**Table 1:** Genes observed in all individuals until or after a particular time point

|  |  |  |  |  |  |
| --- | --- | --- | --- | --- | --- |
| **Seen until** | **0h** | **12h** | **24h** | **48h** | **84h** |
| **#genes** | 14 | 9 | 72 | 52 | 11923 |
| **Mean RPKM when** |  |  |  |  |  |
| **seen** | 0.68 | 0.679 | 1.29 | 1.09 | 32.689 |
| **Unseen before** | **0h** | **12h** | **24h** | **48h** | **84h** |
| **#genes** | n/a | 4 | 2 | 19 | 35 |
| **Mean RPKM when** |  |  |  |  |  |
| **seen** | n/a | 1.078 | 2.212 | 2.769 | 3.034 |

### Different transcripts are degraded at different rates

We sought to better understand the nature of transcript degradation in the RNA samples of lower quality. Given our time course study design, we were able to estimate degradation constants for all genes detected as expressed at all five time-points. To do so, we fit a log-normal transform of a simple exponential decay function (see methods) to quantile-normalised RPKM values for each gene that was detected as expressed in all individuals at all time-points. We considered the slope of this function, *k*, to be a proxy for the decay rate of the gene. We then compared this slope to the mean transcript degradation rate across all genes, which, as a result of our quantile normalization approach, is equal to 0 (thus, a value of 0 indicates no change in the relative rank of that transcript’s expression level across time points). If all genes decay at the same rate, then no slopes should significantly differ from the mean value. However, at an FDR threshold of 1%, we found that 7,267 of the 11,923 genes tested (60.95%) were associated with degradation rates that were significantly different from the mean (figure 3). Of these genes, 3,522 had a negative slope (that is, they were degraded significantly faster than the mean degradation rate) and 3,745 had a positive slope (that is, these transcripts were degraded significantly slower than the mean degradation rate).

**Fig 3.**
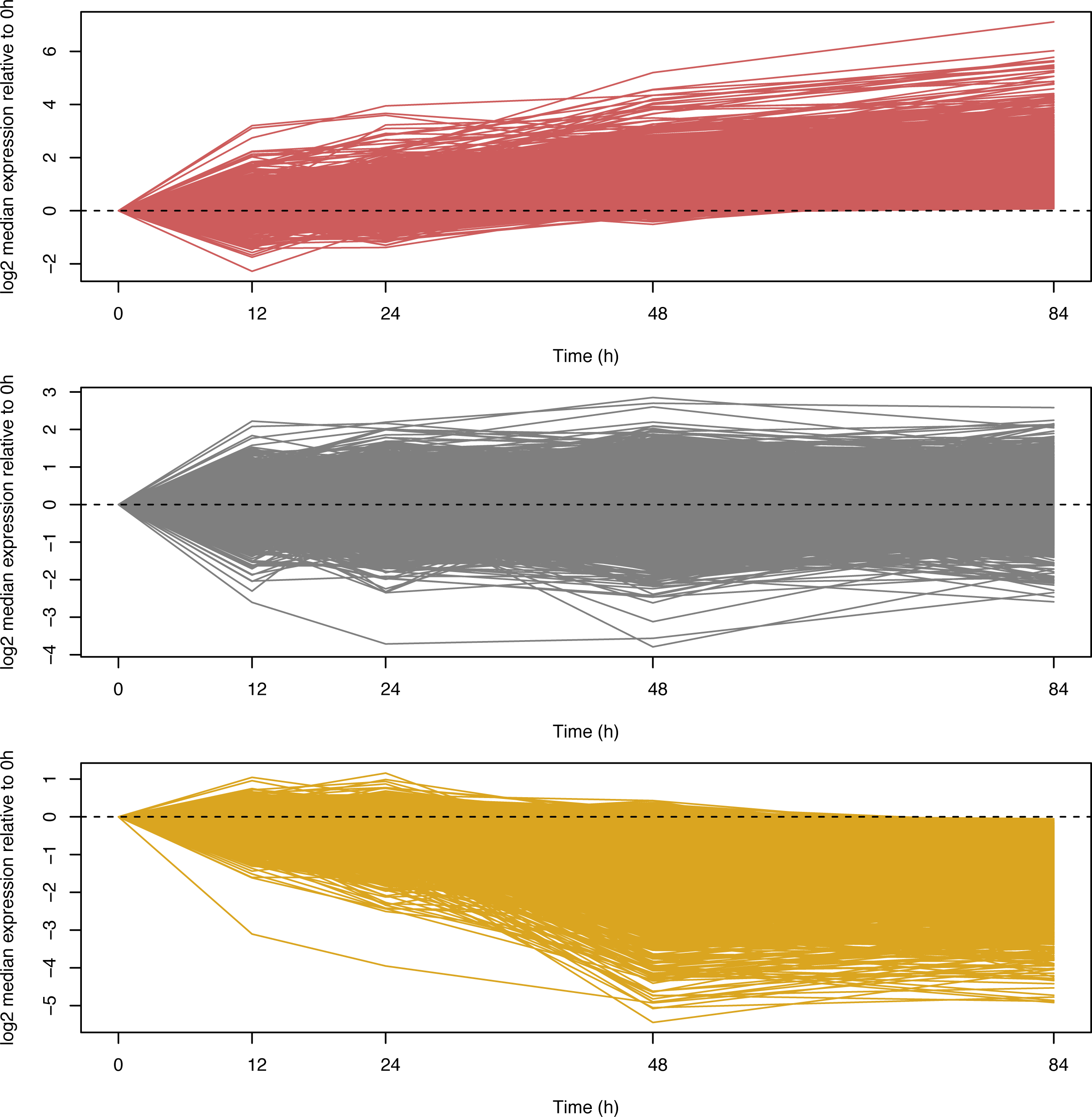
Log_10_ median abundance of genes across all three individuals relative to 0 h. Plots are separated by slope: a. Transcripts with significantly slow rates of degradation relative to the mean rate (identified at 1% FDR, n = 3,745). b. Transcripts that degrade at a rate close to the mean cellular rate (n = 4,656). c. Transcripts with significantly fast rates of degradation relative to the mean rate (identified at 1% FDR, n = 3,522).

Although we might expect RNA degradation in decaying cells to be a random process, GO analysis identified 118 and 293 significantly overrepresented categories amongst slowly and rapidly degraded genes, respectively (FDR = 5%; supplementary tables 3 and 4). A visual inspection of these results reveals that immune terms such as leukocyte activation or immune system process are enriched in genes whose transcripts are degraded faster than average. Terms associated with RNA transcription and protein targeting are overrepresented amongst genes whose transcripts are degraded slower than average. These results are robust to different normalization approaches, to the inclusion of RIN as a covariate in our linear model, or to fitting slopes using RIN instead of time-points. Limiting our analyses to the 1000 bp closest to the 3’ end of transcripts also yields similar results.

We asked what properties, beyond GO functional categories, might be associated with the observed variation in transcript degradation rates. We found that CDS length (p < 10^−12^), %GC content (p < 10^−4^), and 3’UTR length (p < 10^−15^), are all significantly correlated with estimated transcript degradation rate (figure 4A-C), with higher %GC content and increased length of both the 3’ UTR and CDS all associated with faster degradation. However, we found that total transcript length (5’ UTR + CDS + 3’ UTR) is not significantly correlated with degradation rates; instead, intriguingly, targets of both fast and slow degradation have longer transcripts than those that are degraded at an average rate (figure 4D). The correlation between %GC content and CDS length is high (ρ = −0.19, p < 10^−16^), but even when we control for the effects of either variable, the individual effects remain significant predictors of degradation rates (p < 10^−7^). Our data thus suggest that both CDS length and %GC content affect degradation rate, and that observed degradation rates result from complex interactions between multiple forces.

**Fig 4.**
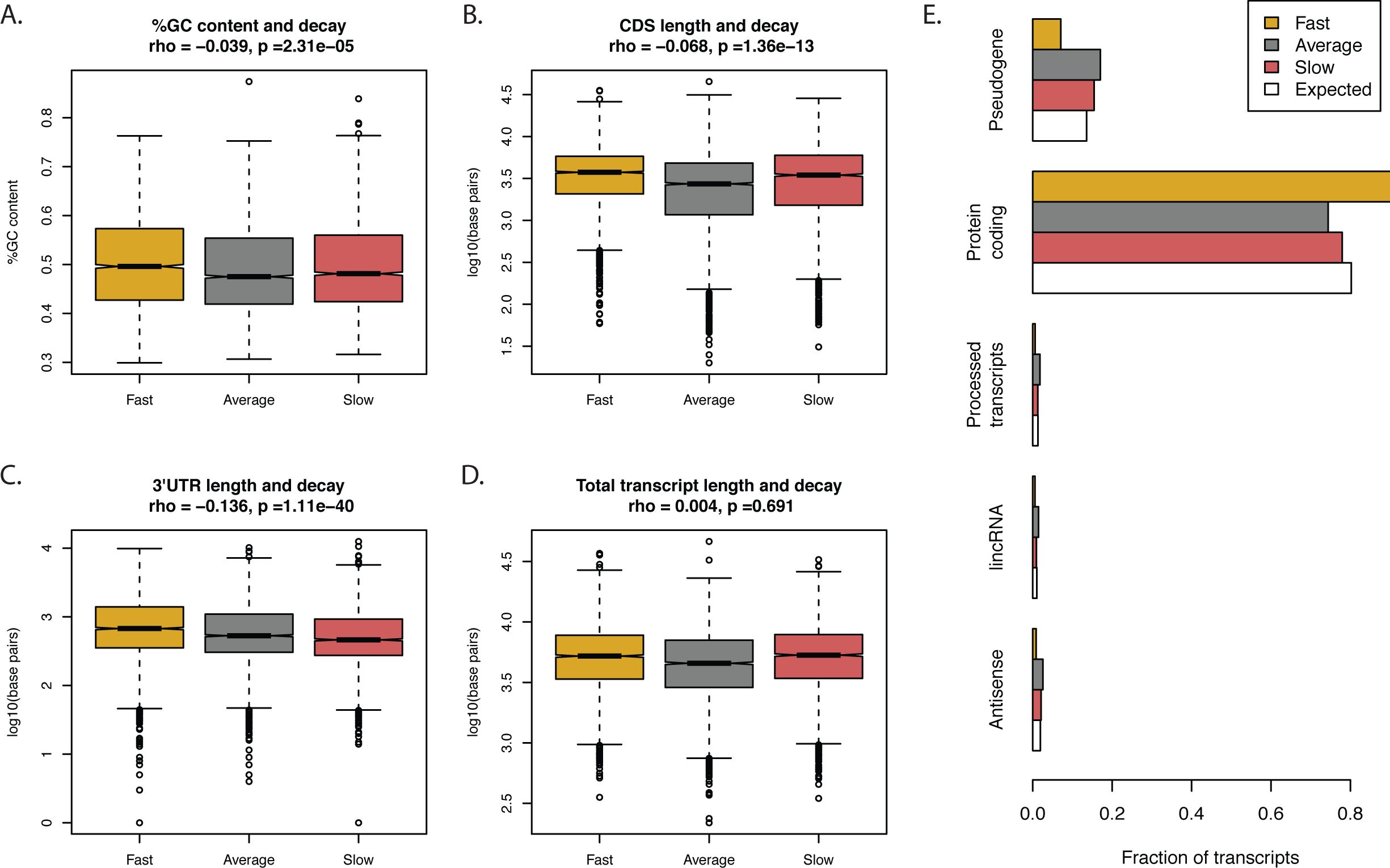
Characteristics of rapidly and slowly degraded transcripts. In all plots, rapidly degraded transcripts are plotted in gold, transcripts degraded at an average rate are plotted in grey and slowly degraded transcripts are in red, a. By transcript %GC content. b. By coding region length. c. By 3’UTR length. d. By complete transcript length. e. By ENSEMBL biotype.

We also sought to investigate whether targets of slow, fast, or average degradation differ meaningfully in terms of broad biological categories. As expected given our poly-A enrichment strategy, most transcripts in our data originate from intact protein-coding genes, but we also observed four other biotypes represented by more than 100 distinct transcripts. The distribution of these biotypes across rapidly and slowly degraded transcripts is not random, with a significant enrichment of pseudogenes among transcripts that degrade slowly (p = 0.015), and an enrichment of intact protein-coding genes among the rapidly degraded transcripts (p < 10^−16^, figure 4E).

### The effect of RNA degradation on the analysis of differential expression

Ultimately, the goal of most RNA sequencing studies is to estimate variation in gene expression levels or to identify genes that are differentially expressed between conditions, individuals, or states. We thus considered the effects of RNA quality on measures of relative gene expression levels between time-points and on estimates of inter-individual variation in gene expression.

As a first step we analyzed the normalized expression data using a GLM approach (see methods) to classify genes as differentially expressed between 0 h and any other time-point. We only considered genes with at least one mapped read in all individuals at all time-points (n = 14,094). At an FDR of 5%, we classified 608 (4%) genes as differentially expressed by 12 h. Both the number of differentially expressed genes and the magnitude of expression changes increase drastically along the time-course experiment (table 2). By 84 h, 9,998 genes (71%) are differentially expressed (FDR = 5%). Roughly half of these genes appear to be more highly expressed in the later time-points than at 0 h. This may seem counterintuitive given that the change in expression is most likely the result of RNA degradation, yet this apparent increase in expression is due to our normalization scheme; indeed, all transcripts in our experiments experience some level of degradation throughout the time course. Post normalization of the data, an apparent elevated expression level in the later time points therefore indicates slow degradation relative to the genome-wide mean rate of RNA decay.

**Table 2:** Number of identified DE genes

GLM: reads ~ time point
| **Time point** | **GLM** | **GLM+RIN** | **Regress RIN, GLM** |
| --- | --- | --- | --- |
| 0h vs 12h | 608 | 5 | 26 |
| 0h vs 24h | 3704 | 5 | 203 |
| 0h vs 48h | 8756 | 47 | 5 |
| 0h vs 84h | 9998 | 42 | 0 |

GLM: reads ~ individual b

| **Individuals** | **GLM** | **GLM+RIN** | **Regress RIN, GLM** |
| --- | --- | --- | --- |
| Ind 1 vs Ind 3 | 69 | 553 | 268 |
| Ind 1 vs Ind 4 | 48 | 401 | 190 |
| Ind 3 vs Ind 4 | 100 | 573 | 299 |

As expected, when we include RIN as a covariate in the model the number of differentially expressed genes across time-points is drastically reduced (fewer than 50 genes are classified as differentially expressed between 0 h and any other time-point; Table 2). These observations confirm that RIN is a robust indicator of degradation levels. Without accounting for RIN, the effect of variation in RNA quality on our data is overwhelming. However, the important question is whether by taking RIN into account we can more clearly identify other sources of variation in the data.

To test this possibility, we focused on variation in gene expression levels between individuals. Without accounting for RIN we classified few genes (48–100; table 2) as differentially expressed between pairs of individuals. This property of the data is also captured by a heat map of sample pairwise correlation calculated using only the top 10% (1,410) most variable genes across individuals at 0 h. As can be seen in figure 5a, while at the early time-points inter-individual differences are the predominant source of variation in the data, degradation overwhelms these differences in the low quality (low RIN) RNA samples from 48 h and 84 h. Hence, inclusion of these time points in our GLM, which considers samples from the same individual but different time points as ‘technical replicates’, obscures much of the true signal of inter-individual variability.

**Figure 5.**
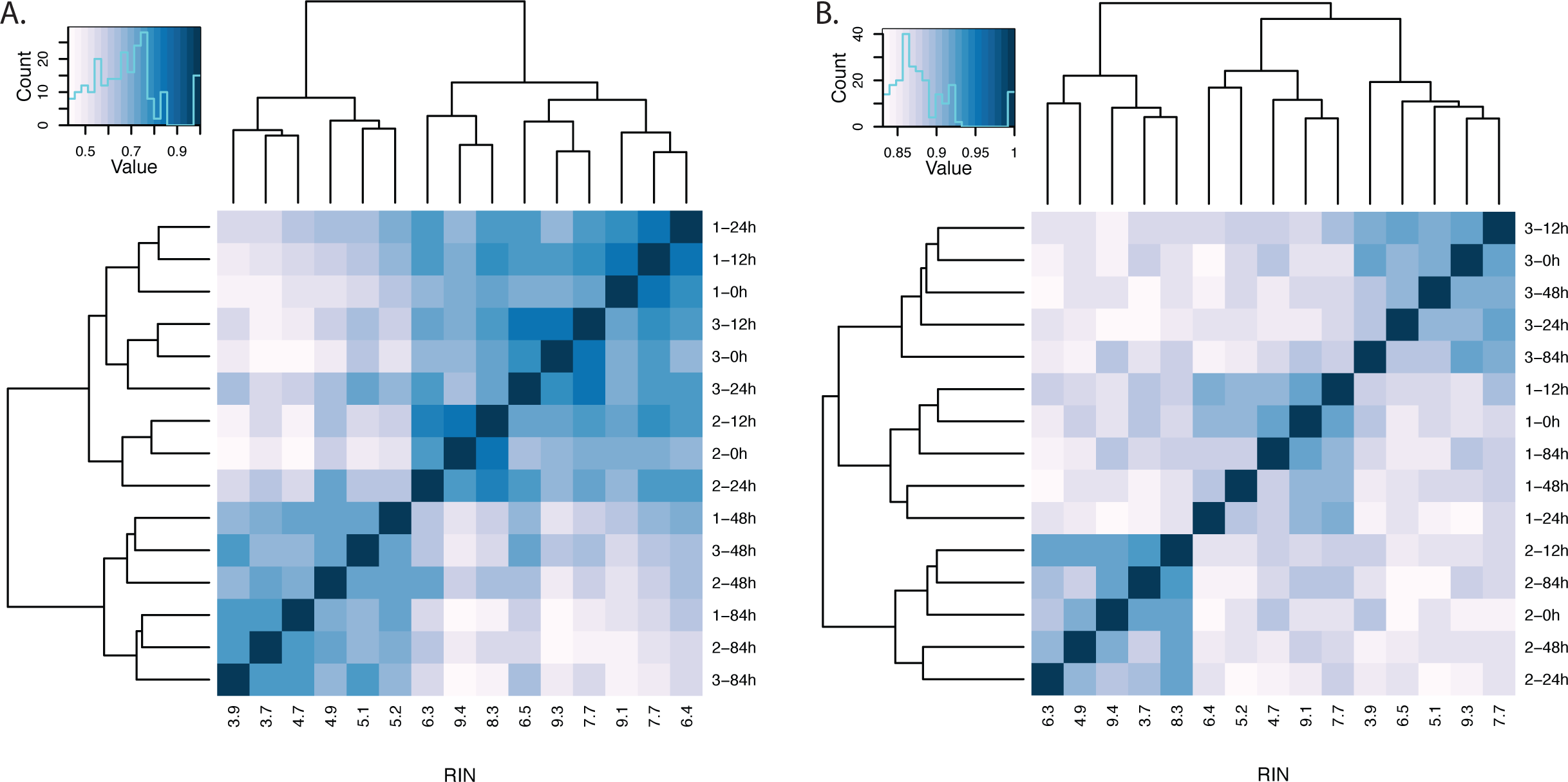
Spearman correlation matrices of the top 10% genes with high inter-individual variance at 0 h. a. Before RIN correction. b. After regressing the effects of RIN.

To recover this signal, we tested two approaches for explicitly accounting for RIN when estimating differential gene expression across individuals: (i) incorporating RIN as a covariate in our GLM; and (ii) analyzing the residuals of gene expression levels after first regressing out RIN from the normalized gene expression data (table 2). Both approaches result in the identification of many more genes as differentially expressed between individuals (401-573 when incorporating RIN directly into our GLM, 190-299 when testing for differential expression using residuals; table 2). We also repeated the pairwise correlation analysis using the same 1,410 most variable genes identified above, but this time we used the residuals after regressing the effect of RIN from the data. The residuals cluster well by individual throughout the entire time course experiment, regardless of RNA quality (figure 5b).

Finally, we examined the overlap between the subset of the 10% of most variable genes across individuals at 0 h (the 1,410 genes used to generate figure 5) and those identified as differentially expressed across individuals as described above (table 3). Of the two approaches we employed to account for the effect of RIN, testing for differential expression after removing the effects of RIN on the data yielded higher concordance between DE genes and those with high inter-individual variance at 0 h, suggesting it may be the better approach.

**Table 3:** DE Genes across pairs of individuals and overlap with top 10% most variable genes at 0h

| **Test** | **GLM individual** |  | **GLM,**<br>**individual + RIN** |  | **Regress RIN,**<br>**GLM individual** |  |
| --- | --- | --- | --- | --- | --- | --- |
|  | **n DE**<br>**genes** | **% overlap** | **n DE**<br>**genes** | **% overlap** | **n DE**<br>**genes** | **% overlap** |
| Ind 1 vs Ind 3 | 69 | 86.96% | 553 | 45.39% | 268 | 75.00% |
| Ind 1 vs Ind 4 | 48 | 89.58% | 401 | 50.12% | 190 | 78.95% |
| Ind 3 vs Ind 4 | 100 | 87.00% | 573 | 49.21% | 299 | 73.91% |
| All individuals | 160 | 85.00% | 1053 | 42.64% | 521 | 71.98% |

## Discussion

Our observations indicate that the effects of RNA degradation following death or tissue isolation are pervasive and can rapidly obscure inter-individual differences in gene expression. Yet, we also found that by using RNAseq nearly all genes observed at our first time-point could still be detected even in severely degraded RNA samples, but the estimated relative expression levels were drastically affected by degradation. Alhough postmortem RNA degradation is considered a non-regulated process, some of the traditional predictors of regulated RNA decay rates in the cell are associated with variation in RNA quality in our data. For example, longer protein coding regions and 3’ UTRs are correlated with more rapid degradation, similar to previously reported trends [5, 23, 24]. Total transcript length, however, which is a significant predictor of regulated RNA decay in the cell, is not associated with variation in degradation rates in our data.

### The effect of RNA quality on study designs and analysis

We confirmed previous observations of decreasing data quality as time from tissue extraction to RNA isolation increased (supplementary figure 1), both with respect to the number of high quality reads we were able to generate from our sequencing libraries and library complexity. While increased time to RNA extraction did not generally result in the complete loss of transcripts (this only happened in under 8% of cases), the relative expression levels of many transcripts were drastically altered over the time-course experiment, with 61% of genes classified as differentially expressed between 0 h (mean RIN of 9.3) and 84 h (mean RIN of 3.78). This proportion of differentially expressed genes is in line with previous reports of the effects of warm ischemia on human gene expression in tumor biopsies, as assessed using microarrays [16, 18]. The potential of RNA degradation to skew measurements of gene expression levels and obscure biologically meaningful signals is therefore apparent. If there are systematic differences in RNA quality between two classes of samples being compared, we predict that the effect of RNA quality on relative estimates of gene expression levels would be responsible for much of the signal in the data. Furthermore, as degradation rate is to some degree associated with biological function (supplementary tables 3 and 4), it has the potential to confound naïve comparisons of functional annotations as well.

Our observations suggest that some of the effects of transcriptional degradation in *ex vivo* samples cannot be corrected. Library complexity decreases somewhat with lower RNA quality, and some genes (approximately 5%) can no longer be detected at the later time-points. Based on our data we conclude that these effects cannot be corrected by simply sequencing more degraded libraries to a greater depth. On the other hand, the marked effects of RNA degradation on the relative expression level of most genes can be accounted for, to a large degree. Indeed, we found that the inclusion of RIN in our model was sufficient to account for much the effect of degradation and allow us to identify a reasonable number of differentially expressed genes between pairs of individuals in our data. These were not spurious signals generated by our approach; they recapitulate observations made at 0 h (when RNA quality was excellent), but were originally dwarfed by the magnitude of degradation-driven expression changes in the uncorrected data. A similar approach – taking into account variation in RIN – has been previously proposed for the analysis of RTq-PCR data abundance [25].

In a study similar to our own, Opitz *et al.* [26] subjected extracted RNA samples from three advanced human rectal cancer biopsies to degradation through increasingly longer incubation at 60°C and then considered the evidence of time-point/RIN–driven degradation using microarray data. The RIN values spanned by their data mirror values in ours, but the results do not. In contrast to the large RIN-associated effects we observed, Opitz *et al.* reported that of 41,000 tested probe-level data points only 275 demonstrated significant degradation effects, with inter-individual differences being the predominant signal in the data. Assuming that differences in the platforms used (microarrays and RNAseq) are not the reason for this discrepancy, one possible explanation for this stark difference between the studies is that lower RIN scores as a result of degradation of extracted RNA samples (Opitz *et al.)* may reflect substantially different properties than lower RIN scores that are the result of degradation of RNA in decaying cells (our study). Based on the observations of Opitz *et al.* we hypothesize that degradation rates of isolated RNA may be mostly linear and uniform; thus the degradation effects can be accounted for by employing standard normalization approaches. In contrast, degradation rates of RNA in a dying tissue sample, a situation that mirrors more closely conditions likely to be faced by investigators in clinical or field settings, is not uniform across transcripts. Because these differences cannot be neglected in downstream analyses, knowledge of the context in which degradation occurs is therefore crucial.

Our observations suggest that actively mediated degradation of transcripts may occur during necrosis; namely, degradation of RNA in a dying tissue may not be a completely random process. Biologically mediated degradation, whether actively driven by the cell’s decay machinery [1], or simply the consequence of the leakage of RNases into cells as membranes are disrupted, is different from the heat-driven degradation of naked RNA, which in turn is likely to be different from the degradation caused by continued freeze-thaw cycles [27]. It is likely that in a dying tissue, most degradation is initially biologically mediated and directed towards specific classes of transcripts, but as the cellular environment continues to deteriorate, the relative importance of stochastic degradation may increase, giving rise to two simultaneous forces.

Additionally, the increased resolution of RNA sequencing relative to other platforms used to assay gene expression levels [21] is both a hindrance and a boon in this situation, allowing for detection of subtler differences than ever before, but also warranting greater caution when analyzing samples of differing quality.

### Recommendation regarding the inclusion of RNA samples in a study

Previous studies [8–10, 19, 27–29] have sought to provide an RNA degradation threshold below which in-depth analysis of RNA is not recommended. However, these studies have reached conflicting conclusions. Our data suggest that if a simple cut-off value is to be used, a conservative cut-off in the context of RNA degradation in dying tissue samples lies between 7.9 and 6.4, the mean RIN scores associated with 12 h and 24 h in our time course experiment, respectively. We observed few differences in measurements of gene expression between 0 h and 12 h, as evidenced by the low number of genes identified as differentially expressed between the two time-points. Thus, it may be tempting to conclude that so long as all samples in any particular study have roughly similar RINs explicit correction is not necessary. However, when we test for differential expression between other close time-points we identify 3,020 genes as differentially expressed between 48 h and 84 h (difference in mean RIN = 1.3), and 5,293 between 24 h and 48 h (difference in mean RIN = 1). It is clear that measurements of gene expression are extremely sensitive to starting sample quality.

Importantly, in spite of these findings, our observations also indicate that useful data can be collected using RNA sequencing even from highly degraded samples. As long as RIN scores are not associated with the effect of interest in the study (namely, different classes of samples in the study are not associated with different distributions of RIN scores), accounting for RIN scores explicitly can be an effective approach. In our study, we were able to identify differently expressed genes between individuals even when RNA samples with RIN scores around 4 were included. Excluding the samples with RIN values lower than 6.4 in our study would have resulted in a less powerful design than including these samples and globally correcting for RIN values. Given these results, we believe that under most circumstances, the most effective approach may be to include all samples regardless of quality, and explicitly model a measure of RNA quality in the analysis.

## Materials and methods

### RNA degradation

We obtained Buffy coat samples from four adult Caucasian males from Research Blood Components LLC (Boston, MA) and separated PBMCs through a standard Ficoll gradient purification. Each sample was split into aliquots of 4 million live cells and resuspended in 200 uL of PBS. Cells were kept at room temperature and aliquots from each sample lysed every twelve hours by addition of 700 uL of RLT buffer (Qiagen) with beta-mercaptoethanol (Sigma-Aldrich) added at 10 uL BME/1 mL RLT according to manufacturer’s instructions. Lysed cells were immediately frozen, and not thawed until RNA extraction.

### Extraction and sequencing

RNA was extracted using the Qiagen RNeasy kit. Extracted RNA quality was assessed with a BioAnalyzer (Agilent Technologies, Wilmington, DE). From these results we selected five time-points – 0 h, 12 h, 24 h, 48 h and 84 h – that encompassed a large stretch of the degradation spectrum. We then prepared poly-A-enriched RNA sequencing libraries for all 20 individual/time-point combinations according to a previously published protocol [21], using 1.5 µg of total RNA per library in all instances. In all instances, we added 15 ng (1%) of an exogenous RNA spike-in during library preparation, composed of equal parts *Caenorhabditis elegans*, *Drosophila melanogaster* and *Danio rerio* total RNA. Samples were multiplexed and sequenced on four lanes (two per library preparation strategy) of an Illumina HiSeq2000 using standard protocols and reagents. Reads were 50 bp in length.

### Data mapping and normalisation

Data were combined across lanes and data for all libraries were randomly subsampled to the lowest observed number of reads, 12,129,475. Reads were independently mapped to the human (hg19), *D. rerio* (danRer7), *D. melanogaster* (dm3), and *C. elegans* (ce10) genomes using BWA 0.6.2 [30]. All reference genomes were obtained from the UCSC Genome Browser [31]. Only reads that mapped exclusively to a single site in the human genome with one or zero mismatches were retained for downstream analyses. Following mapping, we removed all reads that mapped to more than one genome. At this point we also discarded one individual due to low mappability and read quality.

We calculated RPKM [32] for all ENSEMBL v71 [33] human genes in our data. Genes with multiple transcripts were collapsed into a single transcript containing all exons of the gene; where multiple exons of different size overlapped the same genomic region, the entire region was kept. We discarded all exonic regions transcribed as part of more than one gene. Additionally, we quantile-normalised both RPKM and read count-level data across individuals using the lumiN function in the Bioconductor [34] package *lumi* [35], which controls for, and dampens, technical sampling variance in highly expressed genes. Read counts were log2 transformed prior to quantile normalisation to generate a normal distribution; analyses were carried out on subsequently untransformed counts.

All statistical analyses were carried out using R 2.15.2.

### Calculation of decay rates

We estimated the decay rate of the 11,923 genes with an RPKM > 0.3 in all individuals at all time-points by fitting a first order log-normal transform of the classical first-order decay equation:

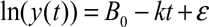

where *y(t)* is the mRNA abundance of a given gene at time *t* (in quantile-normalised RPKM), B_0_ is the abundance at the initial time-point, and *k* the decay rate, with the variance term ε being normally distributed. To control for the high false discovery rate of expressed genes at low expression levels, all RPKM observations < 0.3 were discarded for all subsequent analyses, as in [36].

Length and per-transcript %GC content were calculated using BEDTools (version 2.16.2 [37]), using the same gene models described above. Biotype as well as 5’ and 3’ UTR length were retrieved from ENSEMBL v71. In those instances where there are multiple UTRs associated with the same gene, we used the median UTR length for each gene in all calculations.

### Differential expression and gene enrichment

Differentially expressed genes were identified using the R package *edgeR* [38], utilising a GLM framework with time, individual ID and sample RIN as covariates, as described above. Only those genes with a minimum observation of 1 mapped read across all individuals at all time-points were included. Instead of quantile normalisation as described above, all data were normalised using trimmed mean of M values normalisation (TMM, [39]), which corrects for the observed differences in informative reads between sequencing libraries. Inter-individual variance estimates were generated after variance stabilization of read counts using the *predFC* function in edgeR.

Downstream gene enrichment analyses were carried out using the R package *topGO* [40], using the ‘classic’ algorithm and a minimum node size of 5. All significance values given in the text have been corrected to an FDR of 5% or 1%, using the qvalue method of [41]. In all cases, the background data set included all 14,094 genes with complete observations.

## Acknowledgements

We would like to thank Rachel A. Myers as well as members of the Gilad lab for helpful comments and discussions. This work was supported by NIH grant HG006123 to YG. IGR was supported by a Sir Henry Wellcome Postdoctoral Fellowship. AAP was supported by an American Heart Foundation Predoctoral Fellowship. JT was supported by a Chicago Fellows Postdoctoral Fellowship.

## Supplementary figure legends

Supplementary figure 1. Fraction of reads mapped from generated libraries. All samples were randomly subset to the same depth prior to mapping

Supplementary figure 2. PCA plot of principal components 4 and 5, the only components significantly associated with inter-individual variation in the data. Different colours indicate different time-points, different shapes indicate different individuals.

Supplementary figure 3. a. PCA plot of the 15 samples included in the study based on data from 27,856 genes with at least one mapped read to the 1000-most 3’ base pairs in a single individual. Different shapes indicate different individuals. b. Spearman correlation plot of the 15 samples in the study, using only data trimmed to the 1000-most 3’ bp.

Supplementary figure 4. a. PCA plot of the 15 samples included in the study based on data from 29,156 genes with at least one mapped read in a single individual, after correcting for the effects of RIN on the data. Different shapes indicate different individuals. b. Spearman correlation plot of the 15 samples in the study, after correcting for the effects of RIN on the data.

Supplementary figure 5. Effects of sequencing depth on library complexity. Dashed lines indicate median RPKM in each subset. a-d. Density plots of RPKM values in the 0 h data when subsampled to indicated depths. For comparison, the observed distribution of RPKM values in the 84 h data is plotted in each figure in blue.

